# Bumble bees (Hymenoptera: Apidae) of Ecuador

**DOI:** 10.1101/2025.10.15.682676

**Authors:** Emilia A Moreno-Coellar, Ana B. García-Ruilova, Fernanda Salazar-Buenaño, Pablo A. Menéndez-Guerrero, Esteban Poveda-Proaño, Álvaro Barragán, David A. Donoso

## Abstract

Bumble bee diversity and distribution in the Tropical Andes remain insufficiently documented. Here, we provide the first comprehensive checklist of *Bombus* species for continental Ecuador, integrating data from regional museum collections, published literature, and online biodiversity databases. We further assessed Cytochrome c oxidase subunit I (COI) barcode variation in six species. In total, we documented 15 *Bombus* species occurring across 22 provinces of continental Ecuador. Relative to previous reports, we extend the known elevational ranges of *B. transversalis* and *B. funebris* to higher altitudes, and of *B. excellens* to lower elevations. Three species (*B. mexicanus*, *B. coccineus*, and *B. ephippiatus*) were excluded from the checklist, as earlier records are now considered erroneous. Additionally, citizen science data revealed interactions between eight bumble bee species and 138 plant species, of which 87 species are native. Our findings highlight the value of integrating natural history collections, molecular approaches, and participatory science to secure the long-term conservation of bumble bee diversity in the Tropical Andes.

## Introduction

Bees are one of the most important groups of pollinators worldwide (Buchmann & Nabhan, 1996). Despite bees’ ecological and economic significance, their populations are undergoing alarming declines across the globe (Arbetman et al., 2017; Cameron et al., 2011; Williams & Osborne, 2009). These declines are driven largely by anthropogenic pressures, including habitat loss and fragmentation (Goulson et al., 2015), agricultural intensification (Klein et al., 2007), extensive exposure to pesticides (Gill et al., 2012; Sánchez-Bayo & Wyckhuys, 2019), climate change (Goulson et al., 2015), and the introduction of invasive species, whether accidental or intentional (Morales et al., 2013, 2022). Among them, Bumble bees (Apidae: *Bombus*) are particularly important pollinators in high-altitude and temperate ecosystems due to their efficiency under cold and variable climatic conditions. The genus *Bombus* Latreille, 1802 constitute the monotypic social tribe Bombini, which, along with honey bees(Apini), stingless bees (Meliponini), and orchid bees (Euglossini), forms the group of corbiculates (Cameron, 2007; da Silva et al., 2021). While the greatest diversity of bumble bees is found in temperate regions of Asia, Europe, and South America (Williams, 1998; Michener, 2007), approximately 250 species of bumble bees are currently recognized worldwide. Of these, at least 25 species occur in the Neotropics, and are grouped on seven subgenus (Williams et al. 2008). In Ecuador, updated and integrative information about bumble bee ecology, diversity and distribution is still lacking.

Biodiversity includes not only the number of distinct species but also the interactions among them, which lead to complex ecological networks (Olesen et al., 2007, Hoenle et al., 2019). These interspecific interactions can constrain species distribution and diversity (Bascompte, 2006; Chan et al., 2019), drive evolutionary processes (Ramos & Schiestl, 2019), and play a key role in shaping ecosystem functioning (Garibaldi et al., 2013, Classen et al., 2020). By mapping who interacts with whom, ecological networks reveal not only species associations but also emergent properties at the community level (Vázquez et al., 2009). Among these, plant–pollinator interactions are some of the most extensively studied mutualisms in terrestrial environments (Waser & Ollerton, 2006; Classen et al., 2020). In the neotropics, a vast number of plant species depend on pollination by a diverse assemblage of bees (Velthuis, 1997; Klein et al., 2020), playing a vital role in the functioning of natural, agricultural, and urban ecosystems (Hung et al., 2018; Reilly et al., 2020; Aizen et al., 2022).

Molecular techniques, particularly DNA barcoding, have become essential tools for assessing biodiversity. The use of a standardized fragment of the cytochrome c oxidase I (COI) gene has proven to be one of the most reliable methods for species identification worldwide (Hebert et al., 2003a, b; Janzen et al., 2005). DNA barcoding has shown substantial potential for assessing and resolving biodiversity in groups that are difficult to characterize using traditional taxonomic methods (Köhler, 2007). This approach relies on the delineation of molecular operational taxonomic units (MOTUs), which serve as a practical framework for identifying and interpreting patterns of genetic diversity (Blaxter, 2004; Smith et al., 2005). In bees, DNA barcoding has been successfully applied to a range of taxonomic and ecological challenges, including species discovery and delimitation (Sheffield et al., 2009, 2011; Gibbs, 2009; Gonzales-Vaquero et al., 2016) and the association of castes, such as queens and workers, in eusocial species (Packer et al., 2008).

Study bumble bee diversity provides critical insights into their conservation status, population dynamics and ecological roles, supporting early detection of emerging threats to pollinator communities in Andean ecosystems. In this study, we aim to contribute to bumble bee conservation by 1) providing for the first time a catalogue of bumble bee species in Ecuador, 2) providing for the first time COI barcodes for ecuadorian species and 3) documenting the number of interactions occurring between bumble bees and flowers using citizen science data.

## Materials & Methods

### Bumble bee catalogue and identification

To compile a checklist, we first curated voucher specimen records from the QCAZ invertebrate collection at the Pontificia Universidad Católica del Ecuador (PUCE) and from the dry invertebrate collection of the Museo Ecuatoriano de Ciencias Naturales (MECN) at the National Institute of Biodiversity (INABIO), both museum collections located in Quito, Ecuador. Morphological identifications followed Williams (1998), Michener (2007), Rasmussen (2003) and Abrahamovich (2002, 2004 and 2007). Taxonomic validation of all species followed the Catalog of Bees (Hymenoptera, Apoidea) in the Neotropical Region, according to the guidelines established by Moure and Melo (2007). We extracted specimen records from the Global Biodiversity Information Facility (GBIF; https://www.gbif.org/; only museum records) and from the citizen science platform iNaturalist (https://ecuador.inaturalist.org/; only records previously curated and classified under the ‘research grade’ category). Distribution maps showing the extent of occurrence of each species were created using QGIS (v.3.44).

### Barcoding Bumble bees

We amplified a 658bp fragment of the mitochondrial cytochrome oxidase I (COI) gene from 79 specimens of *Bombus*, previously identified to species level using morphological characteristics. These specimens come from INABIO main insect collection, and from samples from the Global Malaise Project (GMECU, Ecuador). DNA was extracted from the hind leg of the specimens in collaboration with Canadian Centre for DNA Barcoding (CCDB) following the standard protocols of the Biodiversity Institute of Ontario (Wilson, 2012), using the C_LepFolF and C_LepFolR primers. Sequences were sorted into Barcode Index Numbers (BINs; Ratnasingham & Hebert, 2013); a provisional taxonomic framework that assigns unique identifiers to clusters of COI barcode sequences. Each BIN represents a molecular operational taxonomic unit (MOTU) based on sequence divergence. Record details, including geographic data, photographs, and sequences, are available at dataset DS-BOMBUSEC (**dx.doi.org/10.5883/DS-BOMBUSEC**) and stored in the Barcode of Life Data System (BOLD; Ratnasingham and Hebert, 2007).

### Bumble bees and their interacting plants

We compiled data on bumble bee–plant interactions from iNaturalist (https://ecuador.inaturalist.org/), a citizen science initiative that has become highly popular for biodiversity monitoring (Mesaglio & Callaghan, 2021; Beccacece et al., 2025). We only recorded an interaction when a bumble bee was perched in the reproductive structures of a flower and when both, bee and plant, could be identified to the species level.

Bumble bee identifications were based on external morphology comparisons according to the taxonomic keys mentioned before, and plants were identified and categorized by their geographical origin (native or exotic) according to the Plants of the World Online catalogue (POWO 2025) and the new catalogue for continental Ecuador non-native flora (Herrera et al., 2025). Unique interaction records were excluded from the analysis to reduce noise and increase reliability.

We constructed interaction networks between bumble bees and their associated plant species. For each species, we calculated the species degree, defined as the number of species from the opposite trophic level with which it interacts (i.e., the number of plant species visited by a bumble bee or the number of bumble bee species recorded on a plant). All networks were generated in R version 4.3.1 (R Core Team 2022) using the package bipartite v2.18 (Dormann, Gruber, & Fründ 2008).

## Results

### Bumble bee catalogue

We curated a total of 1,794 specimens belonging to 15 species (Table 1; Figures 1–15). Of these, 529 specimens representing 12 species were obtained from the QCAZ invertebrate collection at the Pontificia Universidad Católica del Ecuador (PUCE), and 104 specimens from 6 species originated from the dry invertebrate collection and the Global Malaise Project (GMECU, Ecuador) at the Museo Ecuatoriano de Ciencias Naturales (MECN), National Institute of Biodiversity (INABIO). In addition, we compiled 93 occurrence records from 15 species reported in the published literature, and 310 museum specimen records from 12 species available through the Global Biodiversity Information Facility (GBIF). Finally, we incorporated 753 occurrence records from 8 species documented via the citizen science platform iNaturalist.

**Figure 1.**
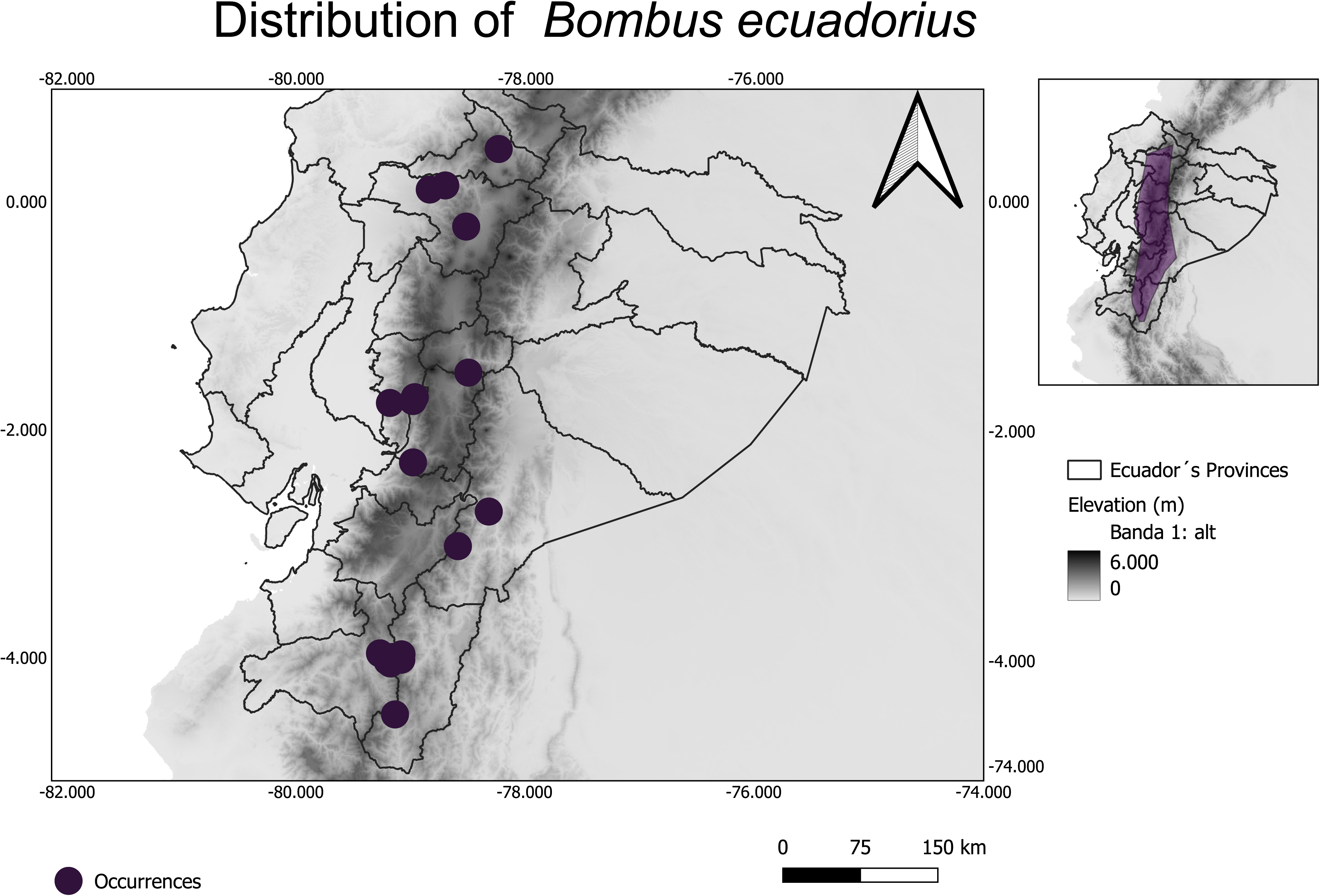
Extent of occurrence of *Bombus ecuadorius* in continental Ecuador.

**Figure 2.**
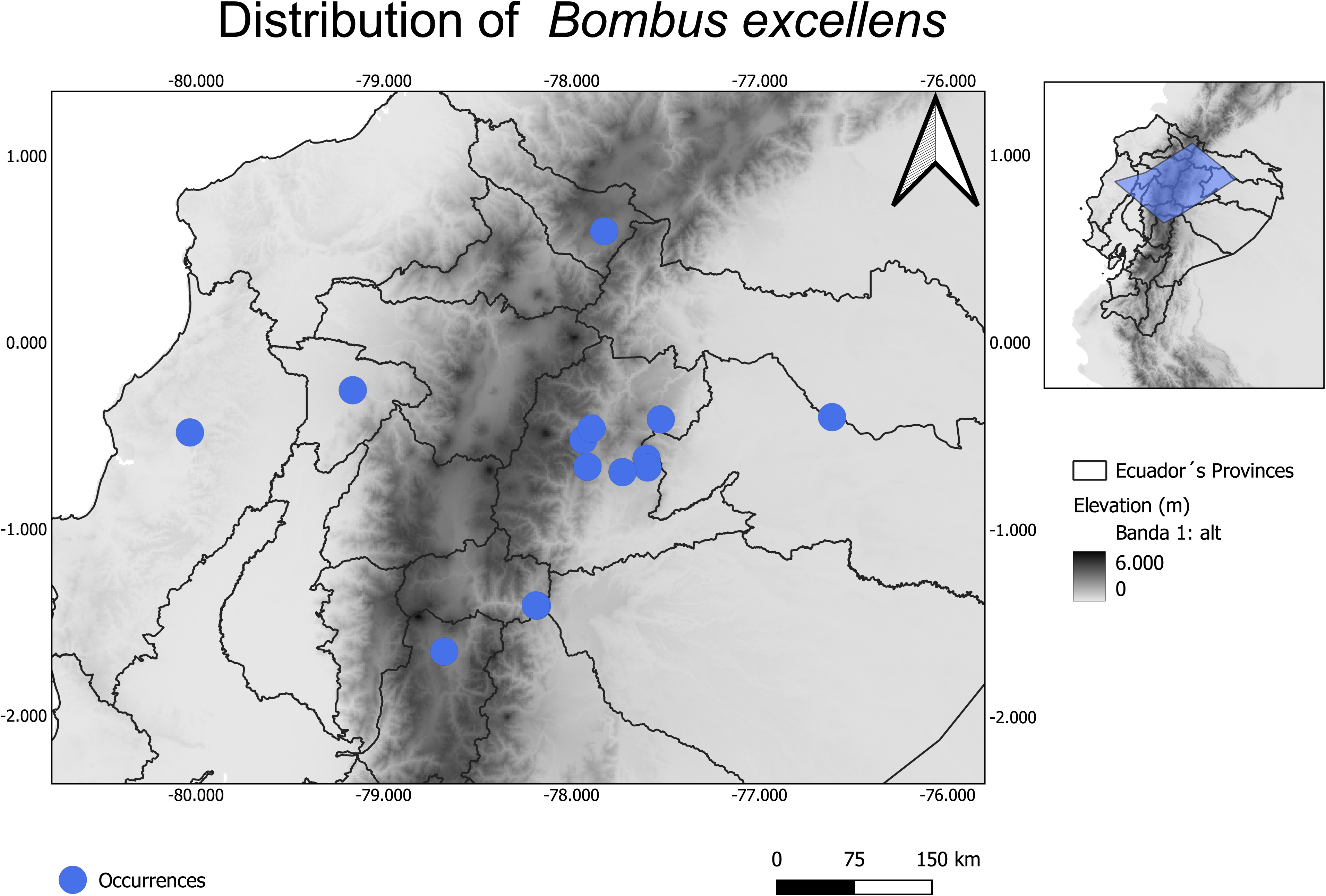
Extent of occurrence of *Bombus excellens* in continental Ecuador.

**Figure 3.**
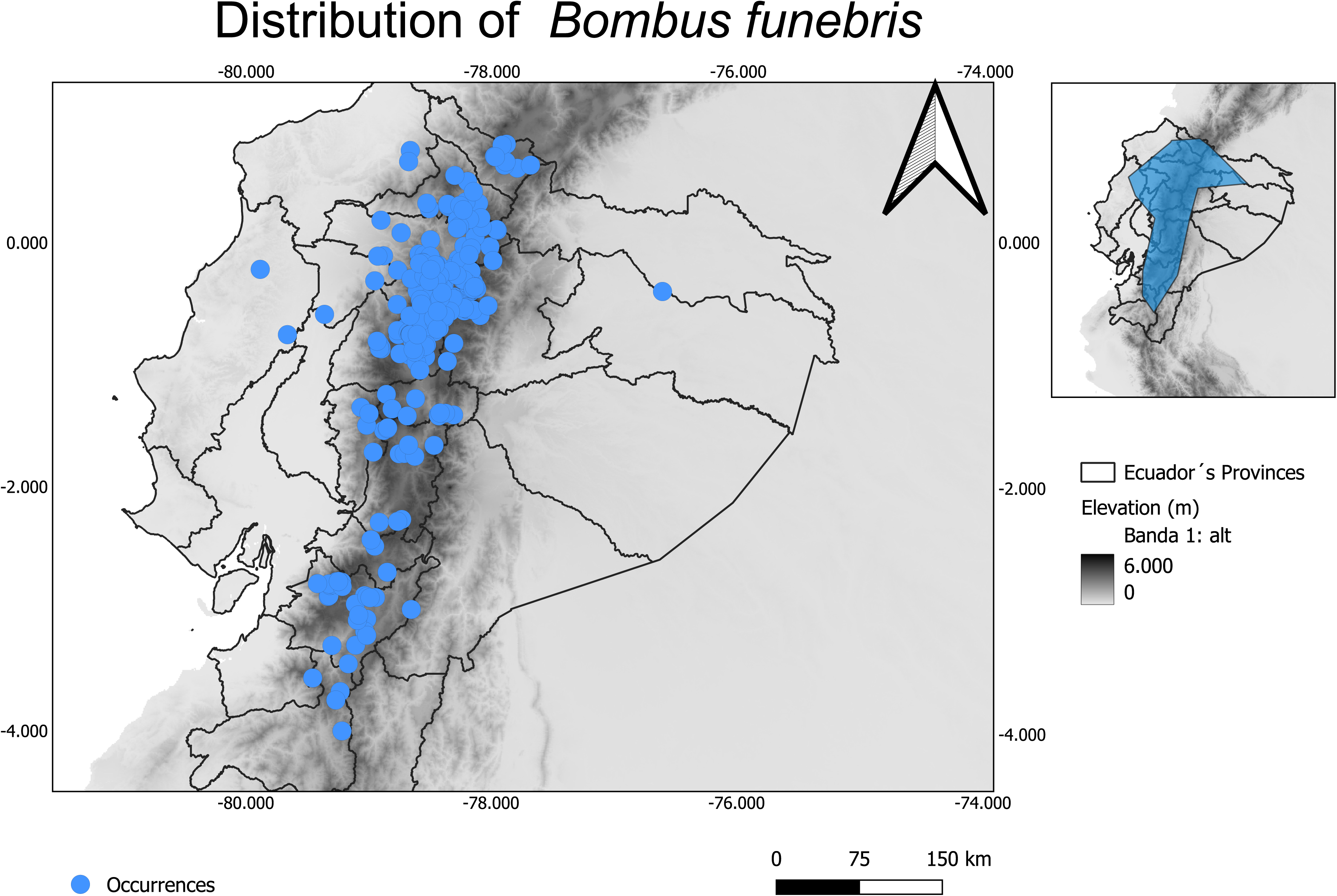
Extent of occurrence of *Bombus funebris* in continental Ecuador.

**Figure 4.**
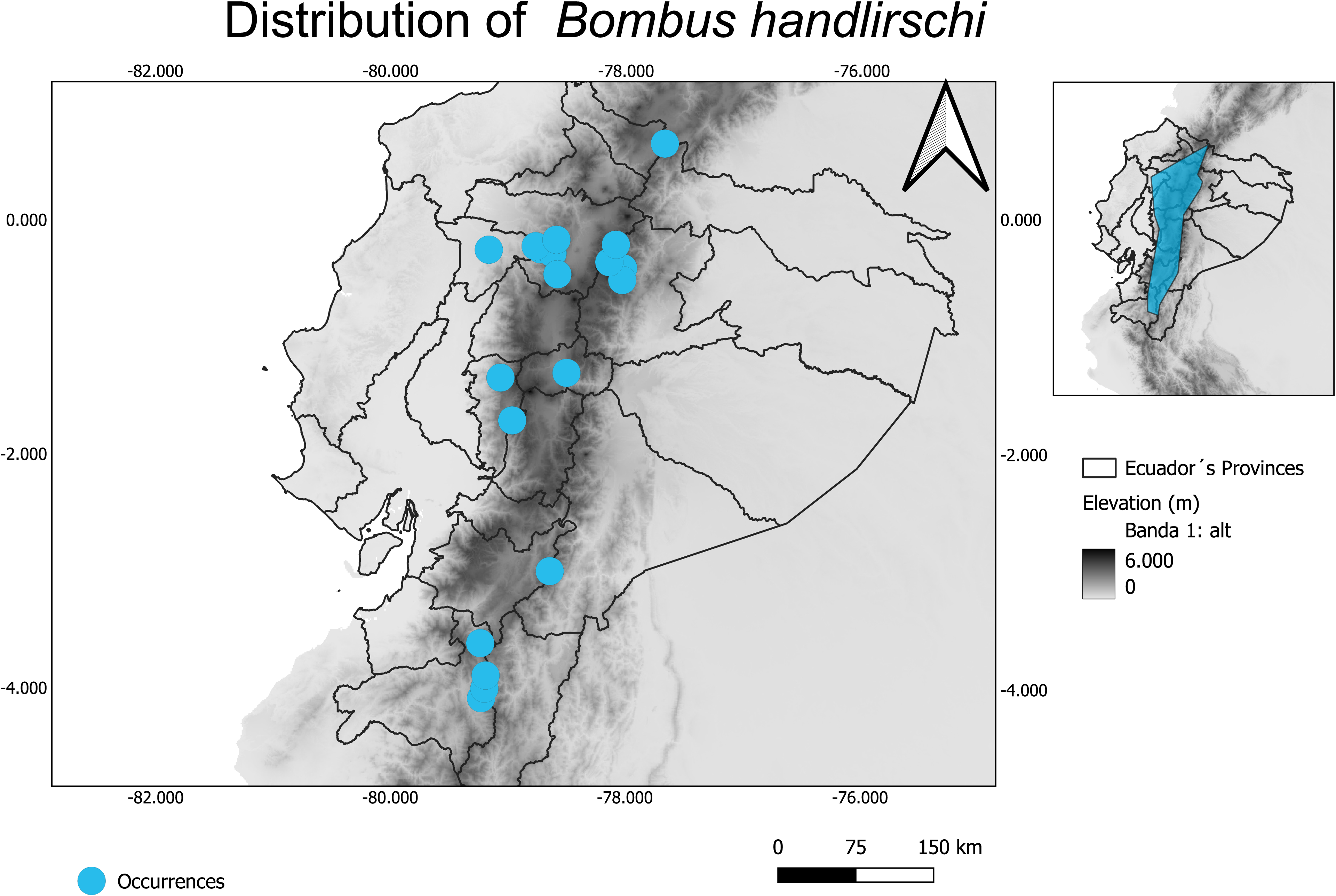
Extent of occurrence of *Bombus handlirschi* in continental Ecuador.

**Figure 5.**
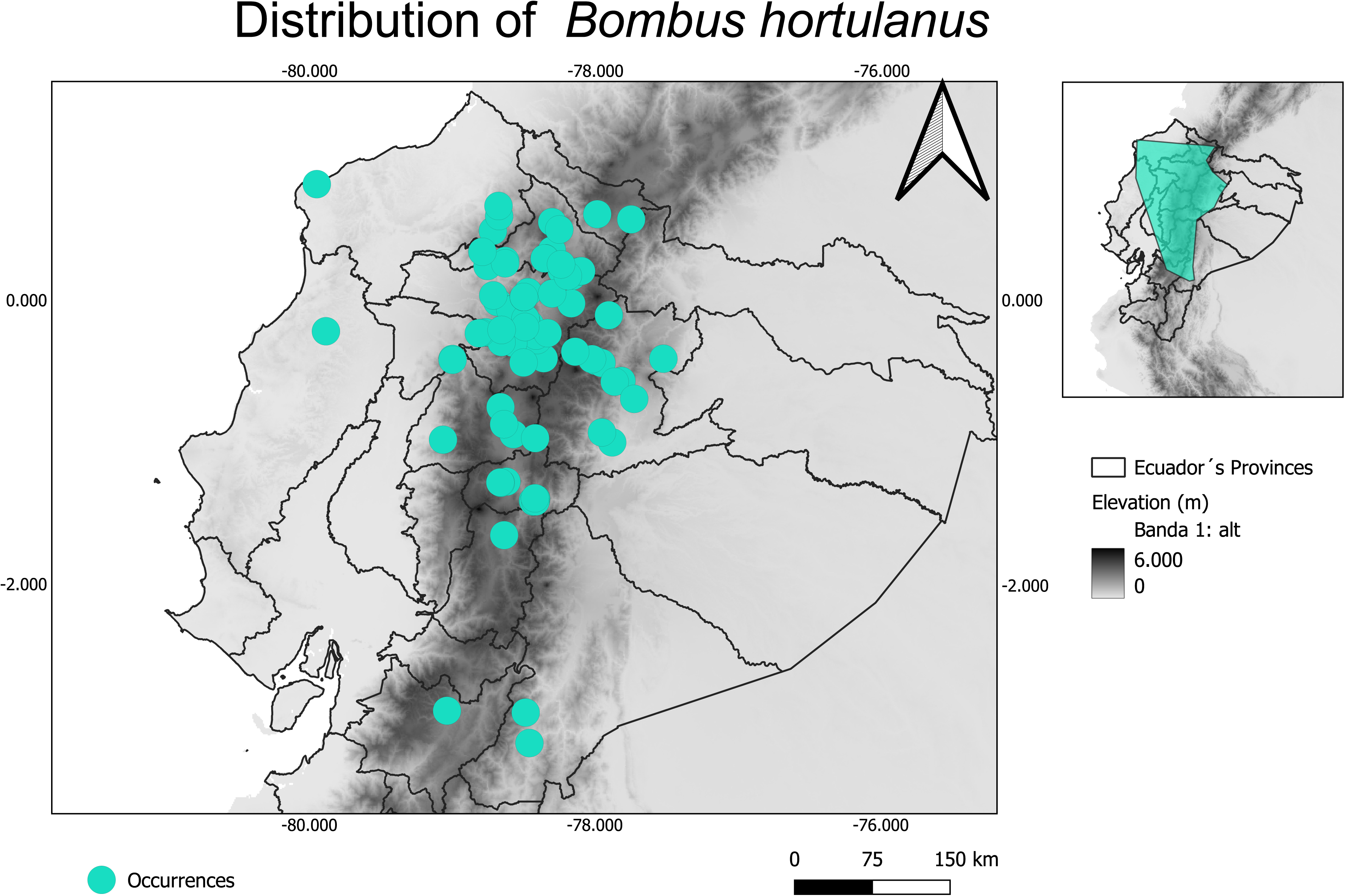
Extent of occurrence of *Bombus hortulanus* in continental Ecuador.

**Figure 6.**
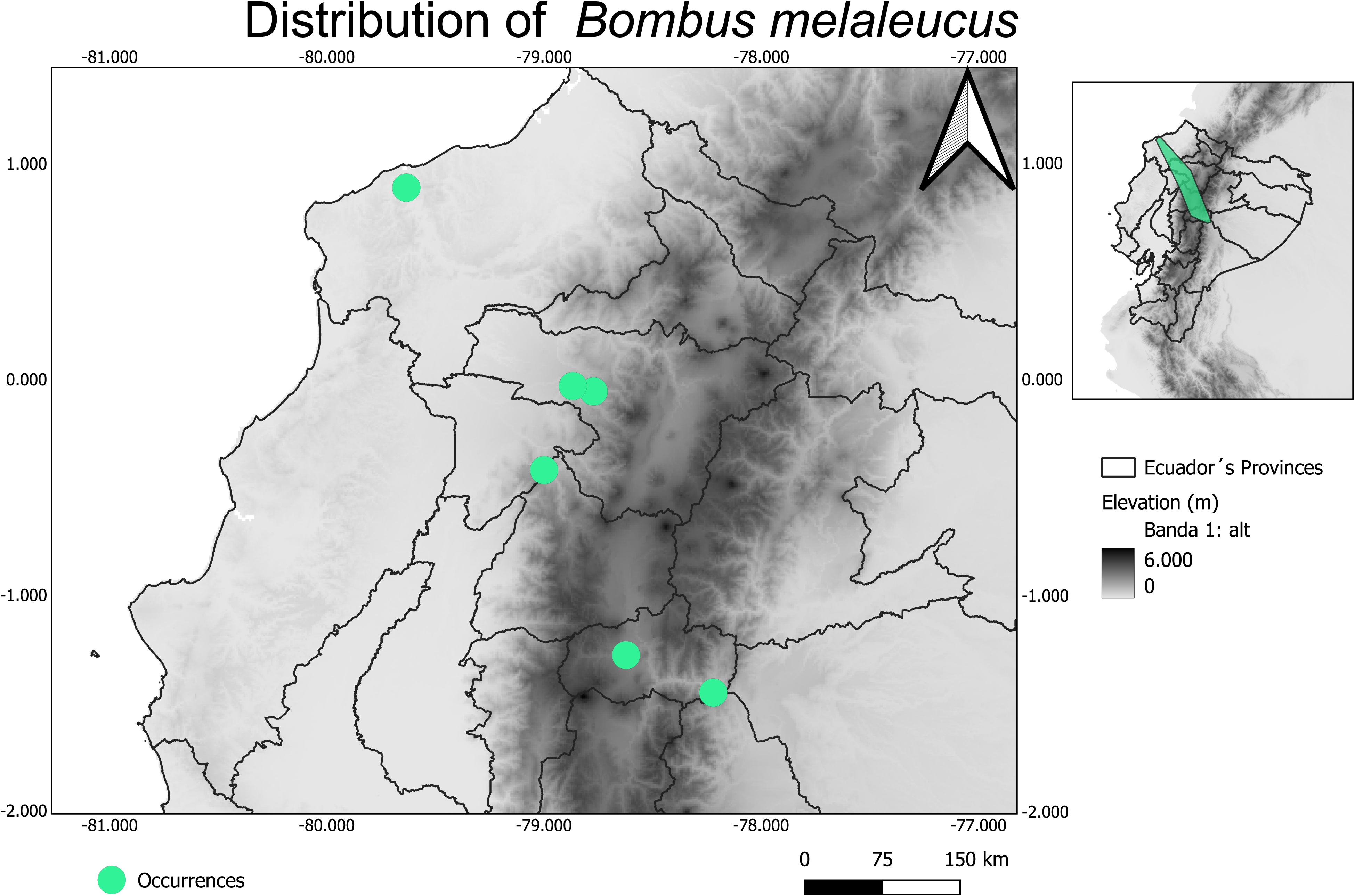
Extent of occurrence of *Bombus melaleucus* in continental Ecuador.

**Figure 7.**
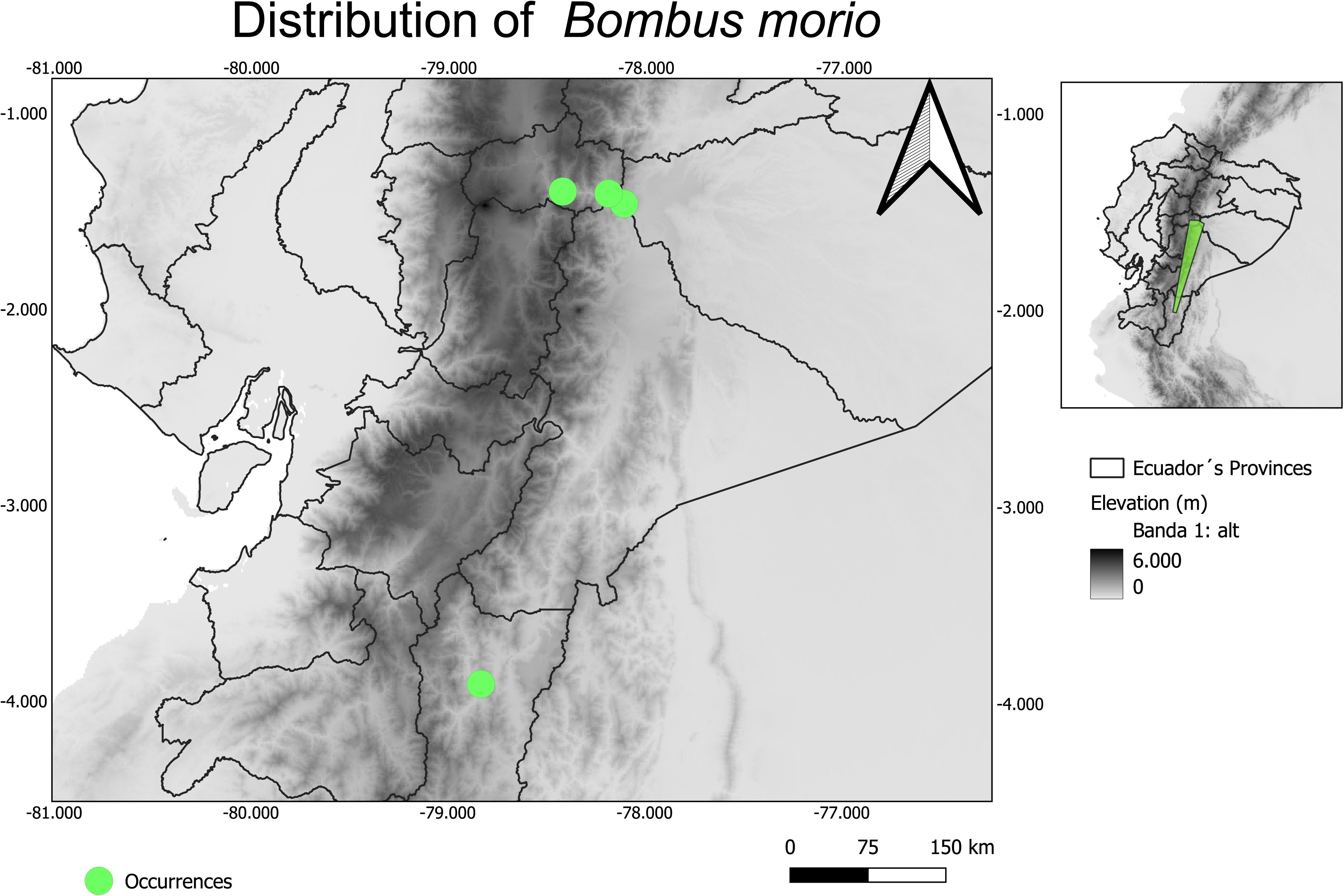
Extent of occurrence of *Bombus morio* in continental Ecuador.

**Figure 8.**
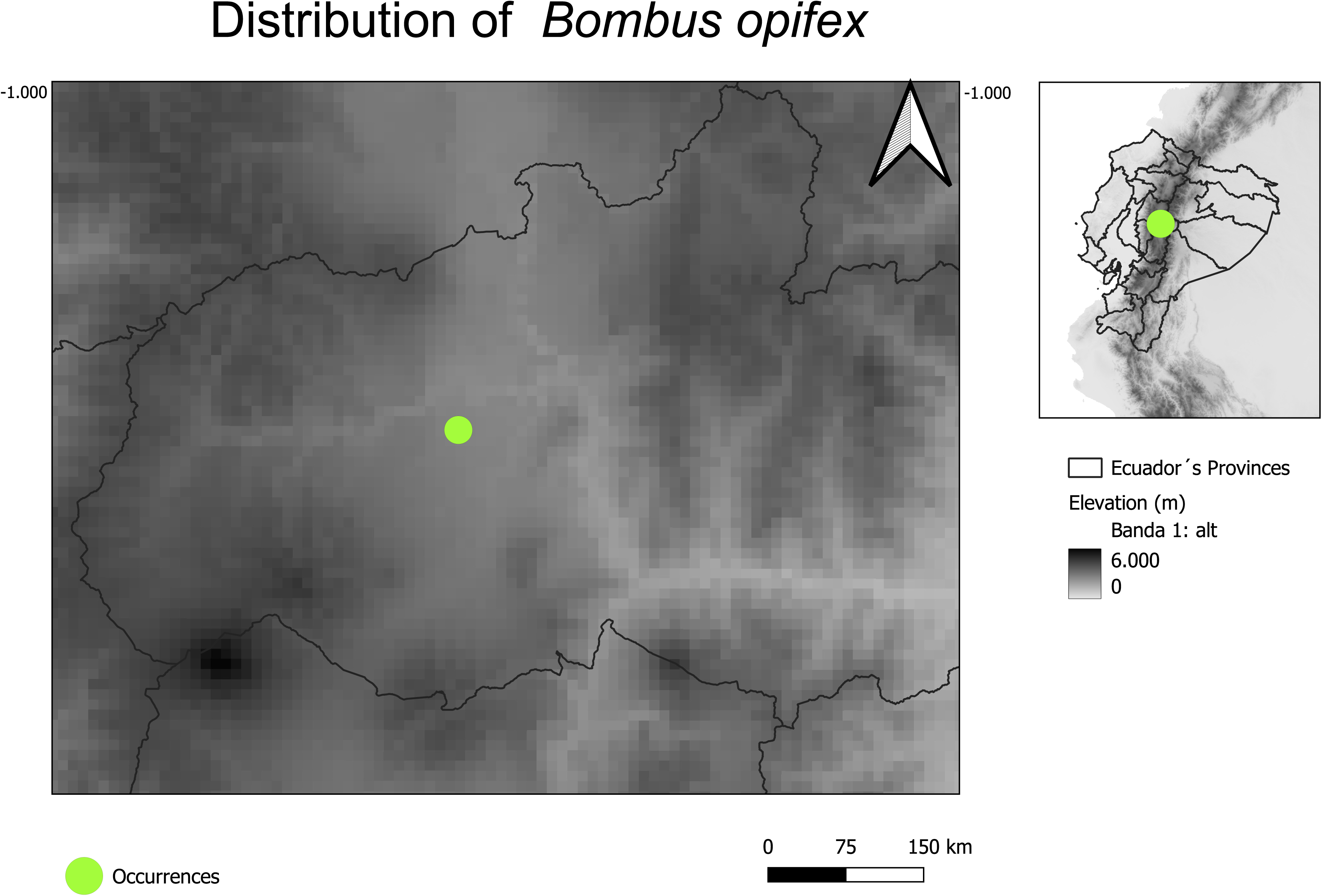
Extent of occurrence of *Bombus opifex* in continental Ecuador.

**Figure 9.**
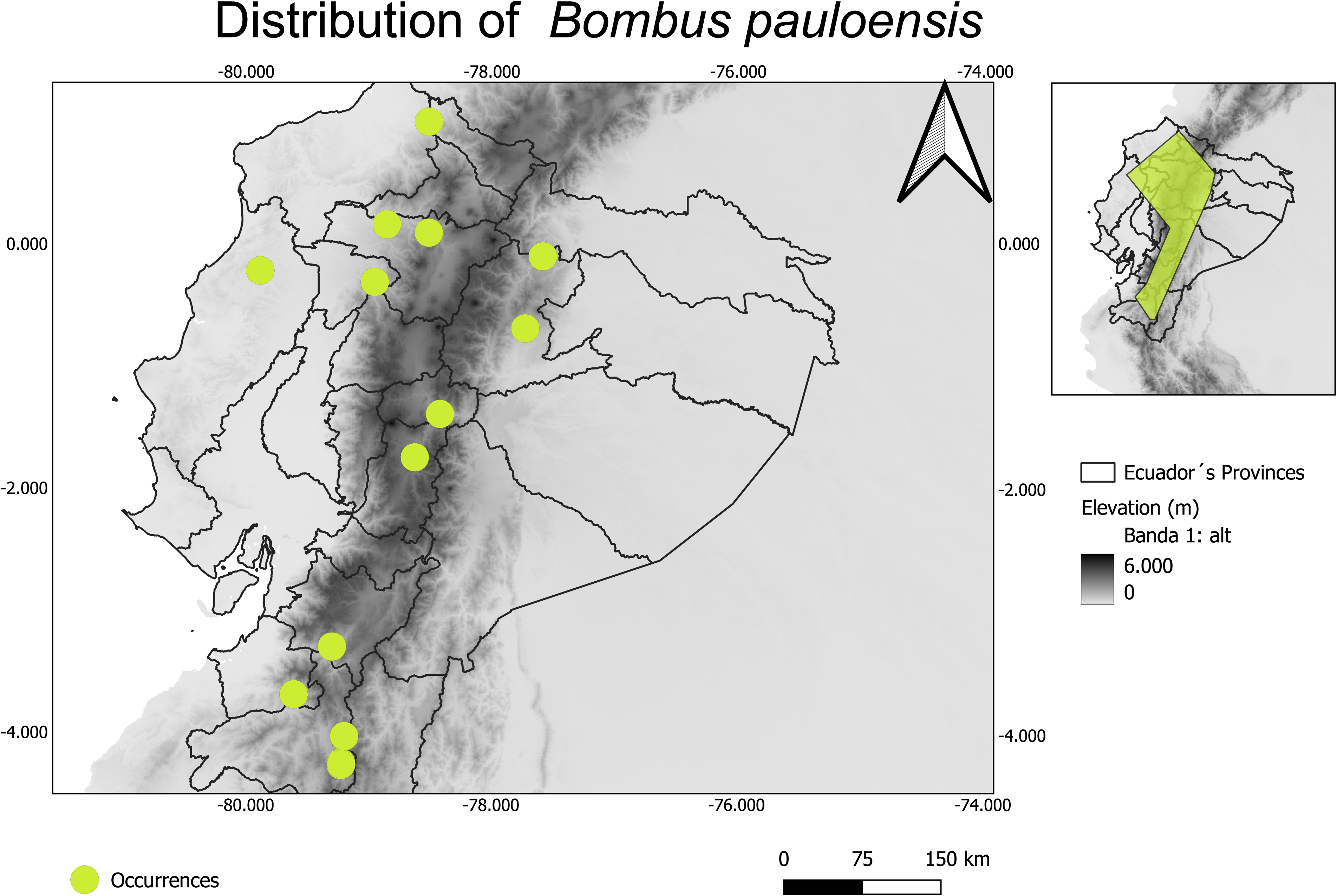
Extent of occurrence of *Bombus pauloensis* in continental Ecuador

**Figure 10.**
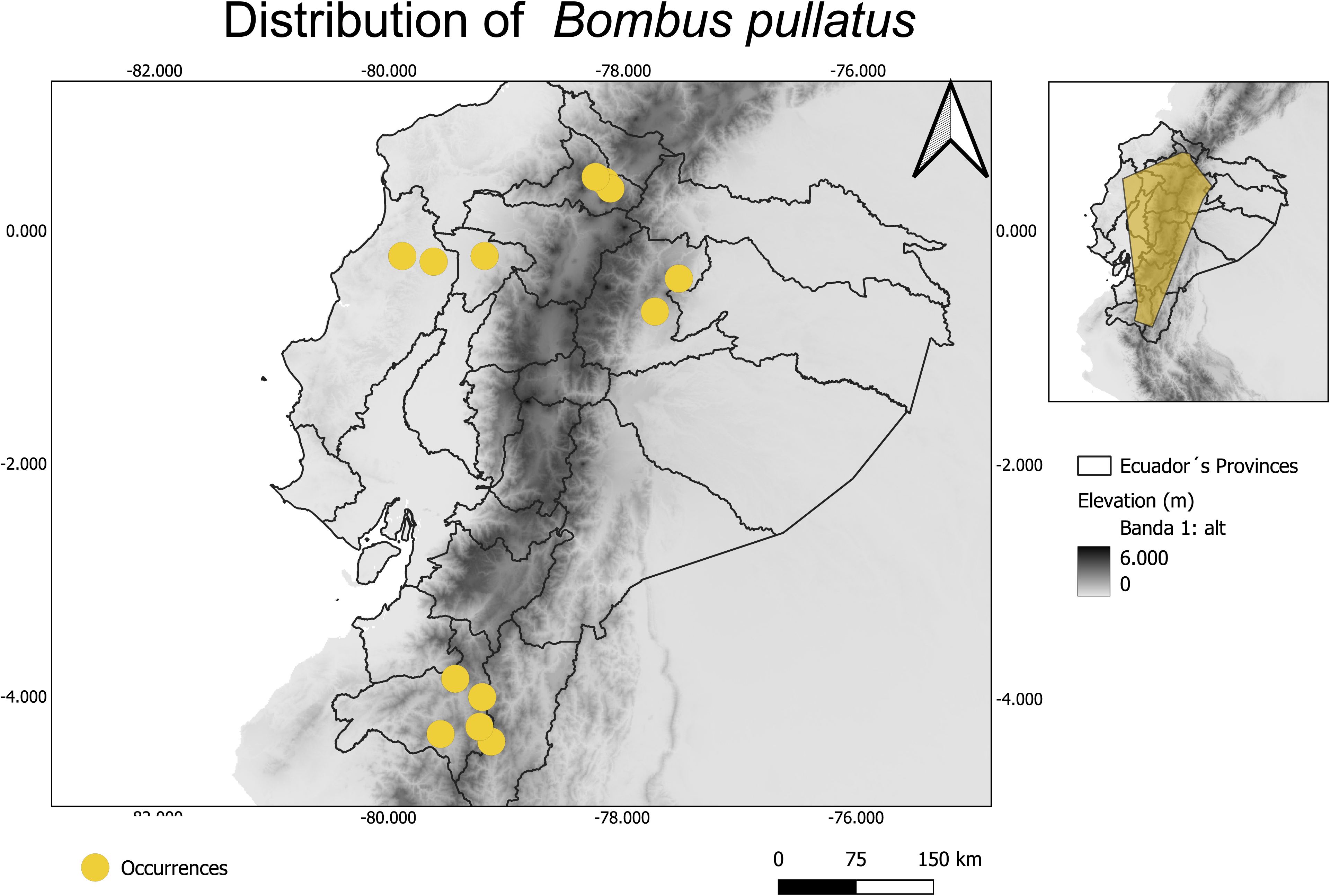
Extent of occurrence of *Bombus pullatus* in continental Ecuador.

**Figure 11.**
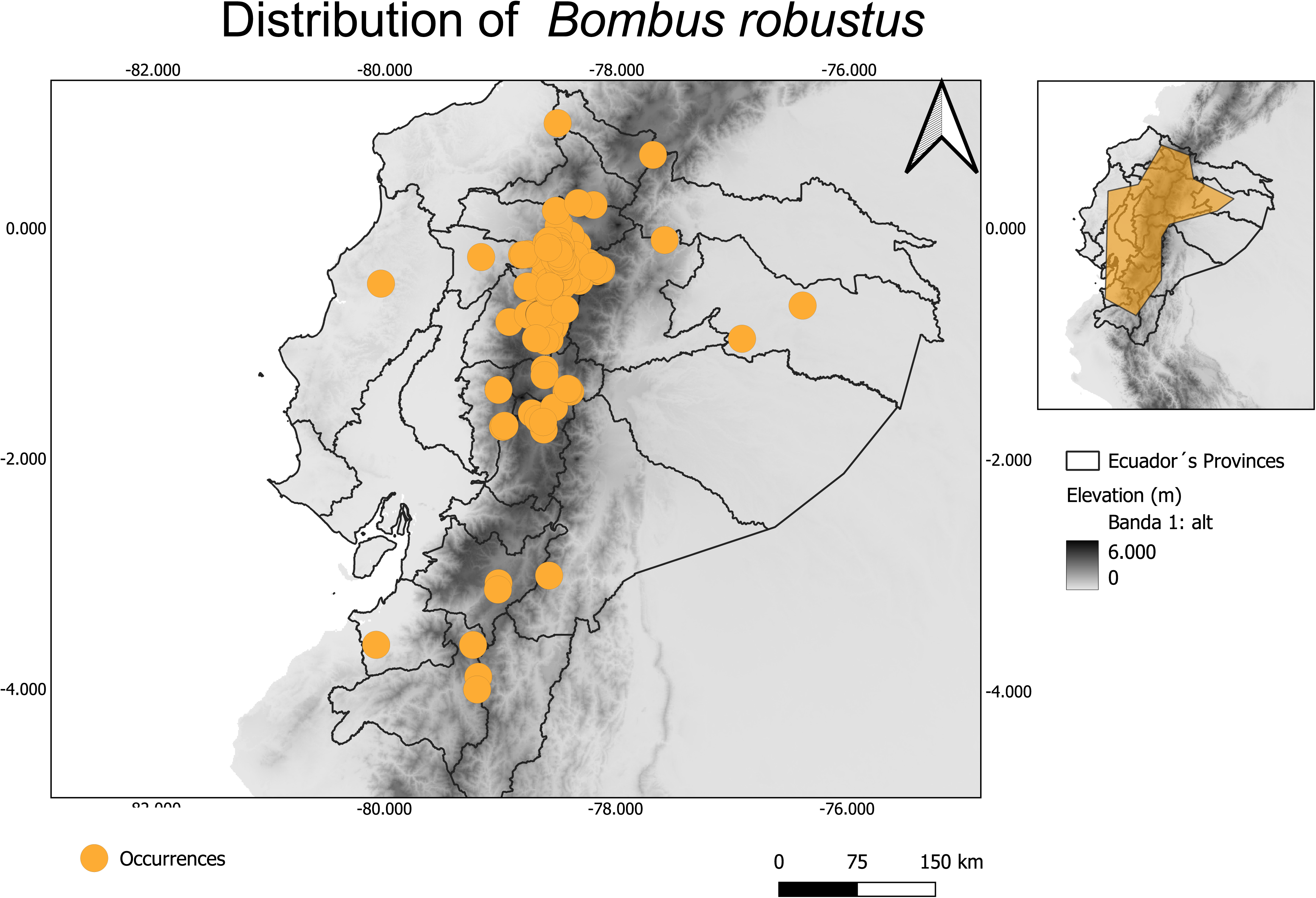
Extent of occurrence of *Bombus robustus* in continental Ecuador.

**Figure 12.**
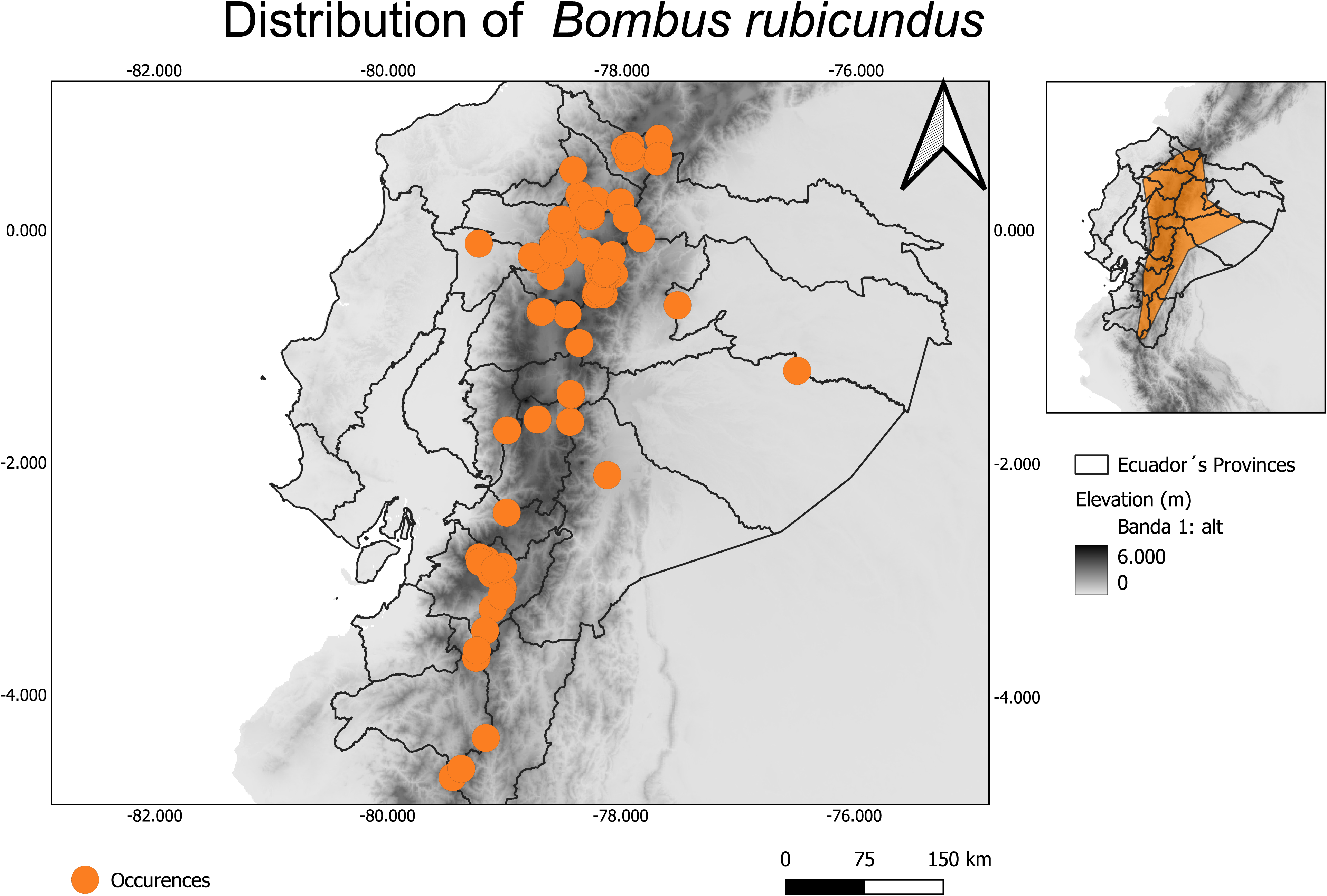
Extent of occurrence of *Bombus rubicundus* in continental Ecuador.

**Figure 13.**
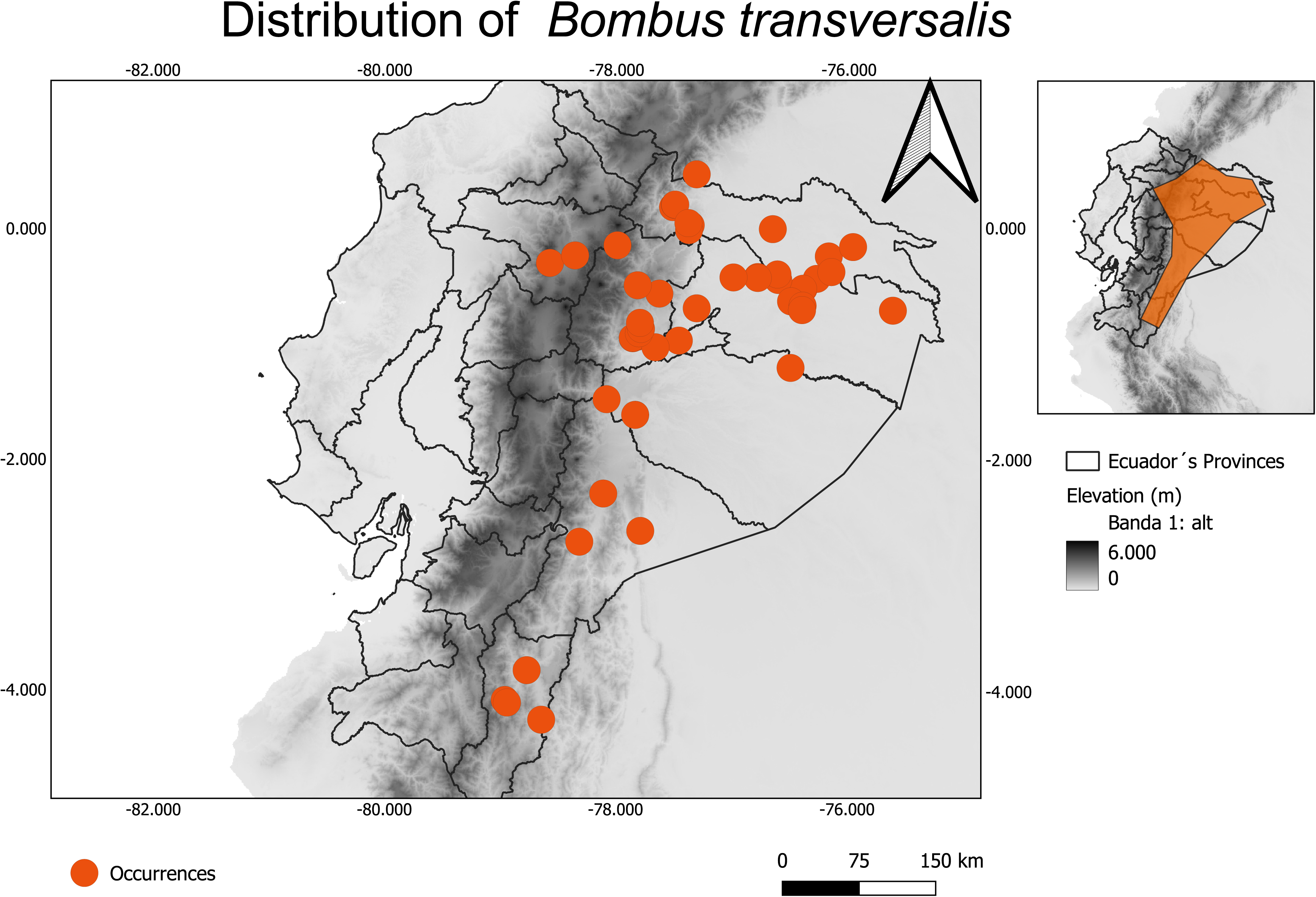
Extent of occurrence of *Bombus transversalis* in continental Ecuador.

**Figure 14.**
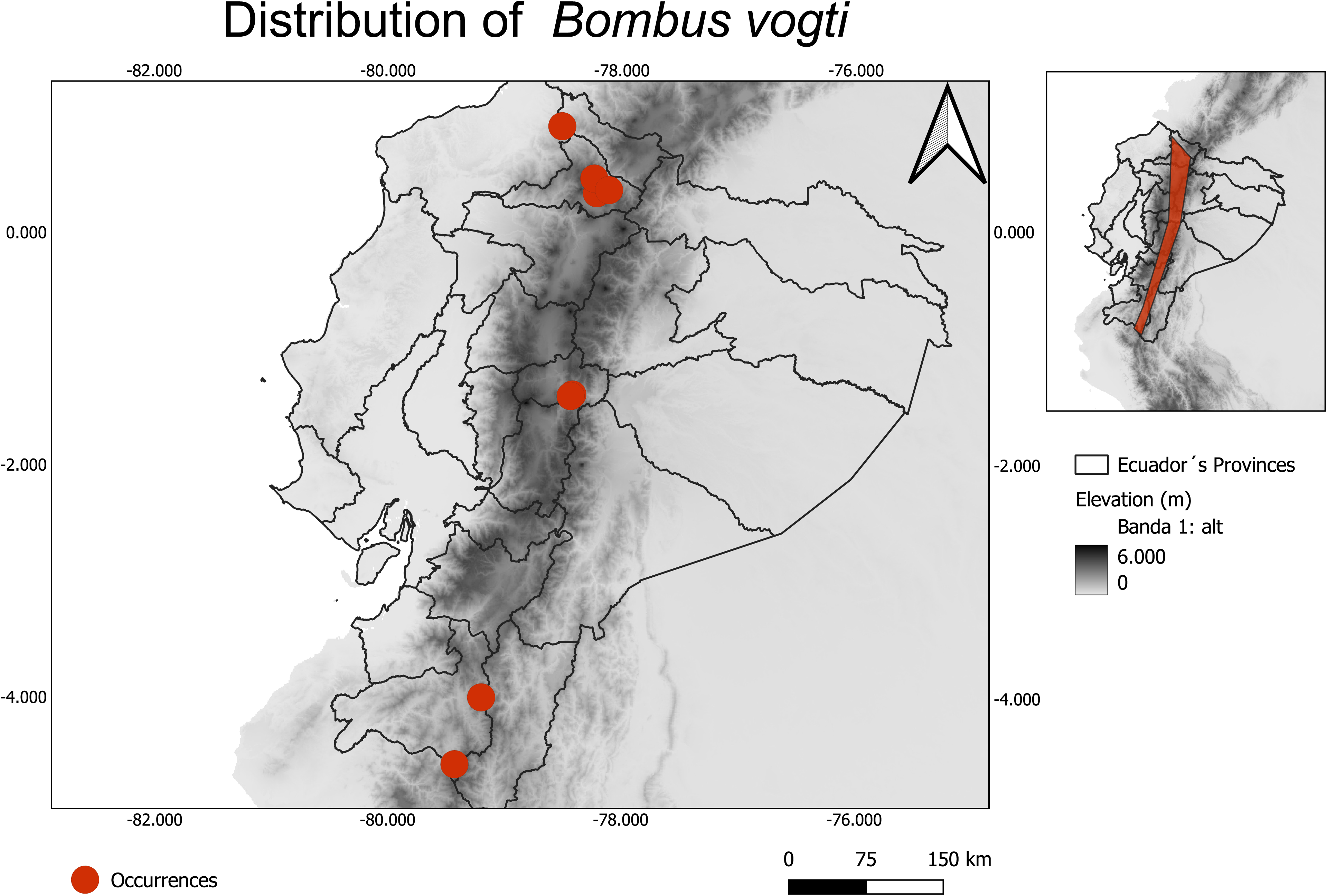
Extent of occurrence of *Bombus vogti* in continental Ecuador.

**Figure 15.**
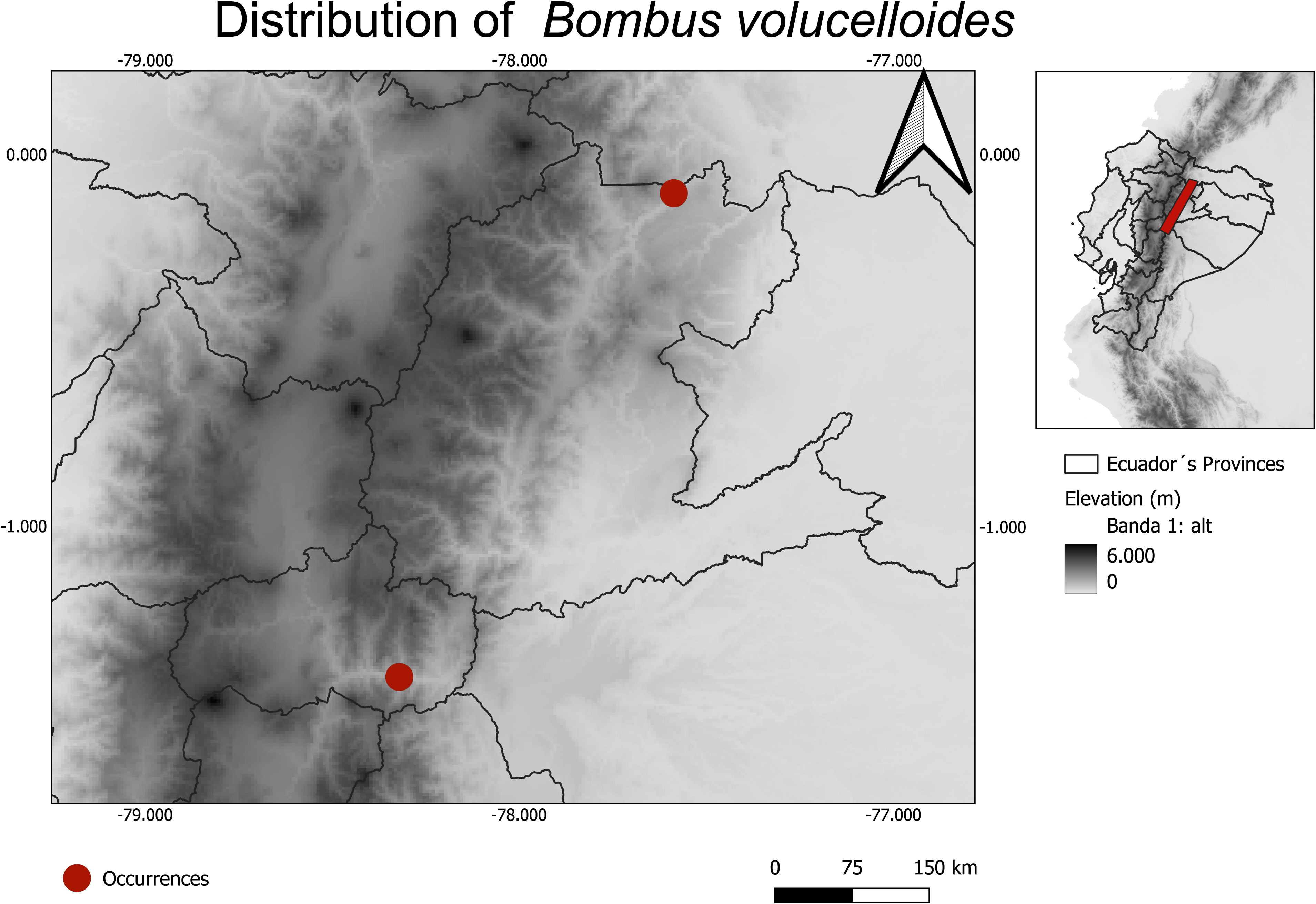
Extent of occurrence of *Bombus volucelloides* in continental Ecuador.

**Table 1.** Checklist of Ecuadorian bumble bees (Apidae: *Bombus*) based on published literature and verified museum records. The list includes valid species names with corresponding taxonomic authorities and the number of examined specimens per species.

| *Bombus*<br>species | BIN | Ecuador |  |  | South America |  |  |  |  |
| --- | --- | --- | --- | --- | --- | --- | --- | --- | --- |
|  |  | Coast | Highlands | Amazon | Ar | Br | Ch | Col | Par |
| *B. funebris* | AAF6139 |  | X |  |  |  | X | X |  |
| *B. hortulanus* | AAJ7887 |  | X | X |  |  |  | X |  |
|  | AFI1835 |  |  | X |  |  |  |  |  |
| *B. pauloensis* | AAZ3039 |  | X |  | X | X |  | X | X |
| *B. robustus* | AEH8532 |  | X |  |  |  |  | X |  |
| *B. rubicundus* | ABX4127 |  | X | X |  |  |  | X |  |
| *B. transversalis* | ABA4058 |  | X | X |  |  |  | X |  |

Further details on these datasets are provided in Supplementary 1.

#### Genus *Bombus,* Latreille 1802

##### Bombus ecuadorius, Meunier 1890

Synonym *Bombus butteli,* Friese 1903

*Bombus ecuadorius* is distributed in Ecuador and Peru at elevations ranging from 1,220 to 3,050 meters above sea level (Abrahamovich et al., 2004; Rasmussen, 2016; Moure & Melo, 2023), with additional records from Bolivia (Rasmussen, 2016). In Ecuador, occurrences span from 732 to 3,000 m.This Andean bumblebee is considered naturally rare, both in museum collections and field surveys. It primarily inhabits native forests, shrublands, grasslands, and likely high-altitude ecosystems. Specimens have been collected exclusively from forested areas and have not been observed in open or agricultural landscapes, suggesting a strong dependence on conserved Andean forest habitats. Due to their specificity, the species is presumed to be particularly vulnerable to habitat degradation. It is currently listed as Data Deficient by the IUCN Wild Bee Specialist Group (Rasmussen, 2016).

##### Bombus excellens, Smith 1879

*Bombus excellens* is distributed across Colombia, Venezuela, Bolivia, Ecuador, and Peru, occurring at elevations between 900 and 3500 meters above sea level (Abrahamovich et al., 2004; Rasmussen, 2016; Moure & Melo, 2023). Here, we report an altitudinal range expansion to 550 m a.s.l., based on a specimen collected in Santo Domingo de los Colorados, Santo Domingo de los Tsáchilas province. Its natural habitat consists of humid tropical premontane forests on the eastern foothills of the Andes, which serve as a transitional zone between the higher montane forests and the Amazonian lowlands. *B. excellens* is considered rare and is currently classified as Data Deficient by the IUCN Wild Bee Specialist Group (Rasmussen, 2016).

##### Bombus funebris, Smith 1854

*Bombus funebris* was originally described from a specimen collected near Quito city (Pichincha, Ecuador). It is a typical Andean species, widely distributed across Colombia, Ecuador, Peru, Chile, and Bolivia, occurring at elevations ranging from 610 to 3800 meters above sea level (González & Engel, 2004; Abrahamovich et al., 2004; Rasmussen, 2016; Moure & Melo, 2023). Here, we report an altitudinal range expansion to 4643 m a.s.l., based on a specimen collected in Ruco Pichincha volcano, Pichincha province. The species is frequently encountered in both natural and disturbed habitats, and their widespread occurrence across diverse environments suggests a broad ecological tolerance and adaptability to a variety of floral resources. Based on current field observations, museum specimens, and citizen science records, its population is presumed to be stable. It is currently listed as Least Concern by the IUCN Wild Bee Specialist Group (Rasmussen, 2016).

*DNA barcode*. The CO1 sequence (650-670 bp) was assigned the BIN: AAF6139 (Fig. 16). A search of the BOLD BIN database confirmed the taxonomic identity of our specimens. The BIN for this species is distributed in Chile, Colombia and Ecuador (see Table 2).

**Figure 16.**
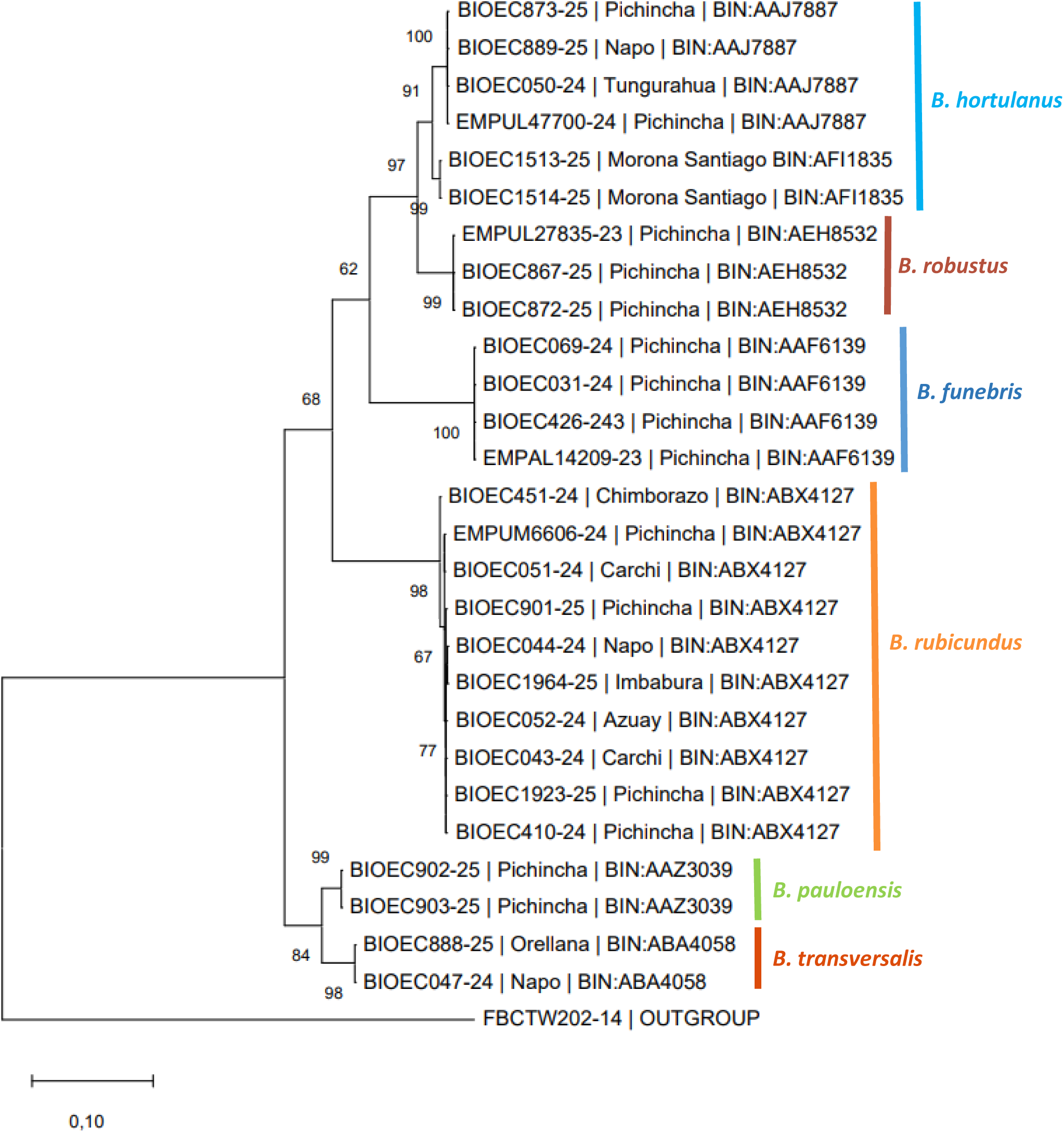
Evolutionary analysis by Maximum Likelihood method of Bombus genus barcodes in Ecuador. The evolutionary history by using the Maximum Likelihood method and Kimura 2-parameter model (Kimura, 1980). The tree with the highest log likelihood(−3.281,61) is shown. The percentage of trees in which the associated taxa clustered together is shown next to the branches. Initial tree(s) for the heuristic search were obtained automatically by applying Neighbor-Join and BioNJ algorithms to a matrix of pairwise distances estimated using the Maximum Composite Likelihood (MCL) approach, and then selecting the topology with superior log likelihood value. A discrete Gamma distribution was used to model evolutionary rate differences among sites (5 categories (+G, parameter = 3.1985)). The rate variation model allowed for some sites to be evolutionarily invariable ([+I], 39.76%sites). The tree is drawn to scale, with branch lengths measured in the number of substitutions per site. This analysis involved 55 nucleotide sequences. Codon positions included were 1st+2nd+3rd+Noncoding. There were a total of 658 positions in the final dataset. The analysis was conducted in MEGA X (Kumar, 2018) utilizing up to 8 parallel computing threads.

**Table 2.** Distribution of *Bombus* BINs in other South American countries. Abbreviations stands for Argentina (Ar), Brazil (Br), Chile (Ch), Colombia (Col) and Paraguay (Par).

| **Bombus species** | **Number of specimens** |
| --- | --- |
| *Bombus ecuadorius* Meunier, 1890 | 5 |
| *Bombus excellens* Smith, 1879 | 2 |
| *Bombus funebris* Smith, 1854 | 223 |
| *Bombus handlirschi* Friese, 1903 | 38 |
| *Bombus hortulanus* Friese, 1904 | 61 |
| *Bombus melaleucus* Handlirsch, 1888 | 1 |
| *Bombus morio* (Swederus, 1787) | 0 |
| *Bombus opifex* Smith, 1879 | 0 |
| *Bombus pauloensis* Friese, 1913 | 11 |
| *Bombus pullatus* Franklin, 1913 | 27 |
| *Bombus robustus* Smith, 1850 | 138 |
| *Bombus rubicundus* Smith, 1854 | 47 |
| *Bombus transversalis* (Olivier, 1789) | 67 |
| *Bombus vogti* Richards, 1933 | 13 |
| *Bombus volucelloides* Gribodo, 1892 | 0 |

##### Bombus handlirschi, Friese 1903

*Bombus handlirschi* is found in Bolivia, Ecuador, Peru, and Venezuela, occurring at elevations between 1830 and 3050 meters above sea level (Abrahamovich et al., 2004; Rasmussen, 2016; Moure & Melo, 2023). In Ecuador, it has been recorded from 550 up to 3970 meters a.s.l. It is currently classified as Least Concern by the IUCN Wild Bee Specialist Group (Rasmussen, 2016).

##### Bombus hortulanus, Friese 1904

The type locality of *Bombus hortulanus* is Baños city (Tungurahua, Ecuador) (Milliron, 1973). It is distributed across Venezuela, Colombia, and Ecuador, occurring at elevations between 760 and 3800 meters above sea level (Abrahamovich et al., 2004; Rasmussen, 2016; Moure & Melo, 2023). In Ecuador, records range from sea level up to 3200 m a.s.l. This species has been reported nesting among grass roots and typically inhabits areas with low to moderate levels of human disturbance (Nates-Parra et al., 2006). It appears to be naturally rare and is currently listed as Data Deficient by the IUCN Wild Bee Specialist Group (Rasmussen, 2016).

*DNA barcode*. The CO1 sequence (650-658 bp) received the assigned BIN:AAJ7887 and AFI1835 (Fig. 16). A search of the BOLD BIN database confirmed the taxonomic identity of our specimens. The BINs for this species is distributed in Colombia and Ecuador (see Table 2).

##### Bombus melaleucus, Handlirsch 1888

*Bombus melaleucus* is distributed across Bolivia, Colombia, Ecuador, Peru, and Venezuela, occurring at elevations between 1524 and 2743 meters above sea level (Abrahamovich et al., 2004; Rasmussen, 2016; Moure & Melo, 2023). In Ecuador, it is found from 200 to 2577 m a.s.l. The species is considered uncommon and is currently classified as Data Deficient by the IUCN Wild Bee Specialist Group (Rasmussen, 2016).

##### *Bombus morio,* (Swederus 1787)

This species is found across Argentina, Bolivia, Brazil, Colombia, Ecuador, Paraguay, Peru, Uruguay, and Venezuela, occurring at elevations from sea level up to 2380 meters above sea level (Abrahamovich et al., 2004; Moure & Melo, 2023; Morales, 2016). In Ecuador, it has been recorded between 918 and 1800 meters a.s.l. It is currently classified as Least Concern by the IUCN Wild Bee Specialist Group (Morales, 2016).

##### Bombus opifex, Smith 1879

This species is distributed across Argentina, Bolivia, Ecuador, Peru, and Paraguay, occurring at elevations between 200 and 4800 meters above sea level (Abrahamovich et al., 2004; Rasmussen, 2016; Moure & Melo, 2023). In Ecuador, it is known from a single historical record from 1956 in Ambato city (Tungurahua, Ecuador), at 2600 meters a.s.l. It is currently classified as Least Concern by the IUCN Wild Bee Specialist Group (Rasmussen, 2016).

##### Bombus pauloensis, Friese 1913

Synonym *B. atratus,* Franklin, 1913

This species is distributed across Colombia, Venezuela, Peru, Ecuador, Bolivia, Brazil, Paraguay, Uruguay, Panama, and Argentina, occurring at elevations ranging from 2800 to 3800 meters above sea level (Abrahamovich et al., 2004; Sasal, 2016; Moure & Melo, 2023; Gonzalez et al., 2004). In Ecuador, it has been recorded from 950 to 2735 m a.s.l. It nests in both rural and urban environments (Nates-Parra et al., 2006; Gonzalez et al., 2004), utilizing a variety of nesting sites from underground cavities (Cameron & Jost, 1998) to aerial nests built in trees (Gonzalez et al., 2004). The species is currently commercially reared for crop pollination in Uruguay (Salvarrey et al., 2013) and Colombia (Cruz et al., 2007; 2008), and is classified as Least Concern by the IUCN Wild Bee Specialist Group (Sasal, 2016).

*DNA barcode*. The CO1 sequence (658 bp) resulted in the BIN:AAZ3039 (Fig. 16). A search of the BOLD BIN database confirmed the taxonomic identity of our specimens.The BIN for this species is widely distributed in South America in countries such as Argentina, Brazil, Colombia, Paraguay and Ecuador (see Table 2).

##### Bombus pullatus, Franklin 1913

*Bombus pullatus* has been recorded in Honduras, Nicaragua, Costa Rica, Panama, Colombia, Venezuela, and Ecuador, at elevations ranging from sea level to 3400 meters (Abrahamovich et al., 2004; Vandame et al., 2015; Moure & Melo, 2023). In Ecuador, it has been found from sea level to 2812 meters a.s.l, inhabiting both conserved and disturbed environments (Nates-Parra et al., 2006). Nesting behavior has been observed both arboreal (Janzen, 1971) and at ground level (Hines et al., 2007), with a notable association with the ant *Acromyrmex octospinosus* (Chavarría, 1996). The species is currently classified as Data Deficient by the IUCN Wild Bee Specialist Group (Vandame et al., 2015).

##### Bombus robustus, Smith 1850

*Bombus robustus* is distributed across Colombia, Venezuela, and Ecuador, occurring at elevations ranging from 762 to 3800 meters above sea level (Moure & Melo, 2023; Abrahamovich et al., 2004). In Ecuador, it has been recorded between 500 and 4000 meters a.s.l. The species is threatened by habitat loss and appears to be naturally rare throughout its range. However, there is limited information on recent population trends, size, and distribution. It is currently classified as Data Deficient by the IUCN Wild Bee Specialist Group (Rasmussen, 2016).

*DNA barcode*. The CO1 sequence (650-658 bp) was assigned the BIN:AEH8532 (Fig. 16). A search of the BOLD BIN database confirmed the taxonomic identity of our specimens. The BIN for this species is distributed in Ecuador and Colombia (see Table 2).

##### Bombus rubicundus, Smith 1854

*Bombus rubicundus* is typically Andean and is distributed across Colombia, Venezuela, Ecuador, and Bolivia, at elevations ranging from 1000 to 4000 meters above sea level (Abrahamovich et al., 2004; Rasmussen, 2016; Moure & Melo, 2023). In Ecuador, it was reported at altitudes between 2200 and 4000 meters a.s.l. Here, we report an altitudinal range expansion from 400 m a.s.l, based on a specimen collected in Moraspungo, Cotopaxi province. There appears to be limited information regarding the population trends and current distribution of this species, and it is currently classified as Data Deficient by the Wild Bee Specialist Group of the IUCN (Rasmussen, 2016).

*DNA barcode.* The CO1 sequence (658 bp) received the assigned BIN:ABX4127 (Fig.16). A search of the BOLD BIN database confirmed the taxonomic identity of our specimens. The BIN for this species is distributed in Ecuador and Colombia (see Table 2).

##### Bombus transversalis, (Olivier 1789)

*Bombus transversalis* is distributed across Colombia, Ecuador, Venezuela, Bolivia, French Guiana, Guyana, Suriname, Brazil, and Peru (Abrahamovich et al., 2004; Sasal, 2016; Moure & Melo, 2023). In Ecuador, it has previously been recorded at elevations ranging from 220 to 1100 meters above sea level. Here, we report an altitudinal range expansion to 1568 m a.s.l., based on a specimen collected in Archidona, Napo province. Colonies of this species can become relatively large. Taylor and Cameron (2003) described its nest architecture in the Amazon regions of Colombia, Brazil, Peru, and Ecuador: nests are typically conical or dome-shaped and are constructed on the surface using leaves, rather than underground. *Bombus transversalis* is currently listed as Least Concern by the IUCN Wild Bee Specialist Group (Sasal, 2016).

*DNA barcode.* The CO1 sequence (658 bp) received the assigned BIN:ABA4058 (Fig.16). A search of the BOLD BIN database confirmed the taxonomic identity of our specimens. The BIN for this species is distributed in Colombia y Ecuador (see Table 2).

##### Bombus vogti, Richards 1933

*Bombus vogti* is distributed across Bolivia, Peru, Colombia, and Ecuador, occurring at elevations between 914 and 2017 meters above sea level (Abrahamovich et al., 2004; Moure & Melo, 2023). In Ecuador, it has been recorded at altitudes ranging from 1680 to 2500 meters a.s.l. The species is naturally rare, and there is limited information regarding its population trends and current distribution. It is currently classified as Data Deficient by the IUCN Wild Bee Specialist Group (Rasmussen, 2016).

##### Bombus volucelloides, Gribodo 1892

This species is distributed across Costa Rica, Panama, Colombia, Ecuador, and Venezuela (Abrahamovich et al., 2004; Moure & Melo, 2023). It prefers warm, humid climates with minimal disturbance (Nates-Parra et al., 2006). The species is currently classified as Least Concern by the IUCN Wild Bee Specialist Group.

### Bombus species not included in the catalogue

We excluded literature records of *Bombus mexicanus* and *B. ephippiatus* reported by Franklin (1913), as their presence in Ecuador appears to be erroneous. According to Moure and Melo (2023) and this study, both species are native to Central America and have not been confirmed in South American localities. Similarly, we did not include *B. coccineus*, originally recorded by Milliron (1973), as the available locality data corresponds to a Peruvian site rather than Ecuadorian territory (Abrahamovich & Díaz, 2002).

### Bumble bees and their interacting plants

From iNaturalist records, we documented eight Bombus species interacting with 140 plant species belonging to 39 botanical families, from where 87 species were native and 51 were exotic (Figures 17-19; Supplementary 1). *Bombus funebris* was the bumble bee with the highest number of links, interacting with 78 different plant species from which 51 were native, followed by *B. robustus*, interacting with 53 plant species from which 21 were native, *B. rubicundus* with 32 plant species from which 26 species were native and, *B. hortulanus* with 20 plant species from which 6 were native. In contrast, *B. transversalis* interacted only with 8 plant species and 7 were native species, *B. ecuadorius* with 4 plant species from which 3 were native, *B. excellens* with 3 plant species, 2 native, and finally *B. melaleucus* with 2 plant species from which one was native.

**Figure 17.**
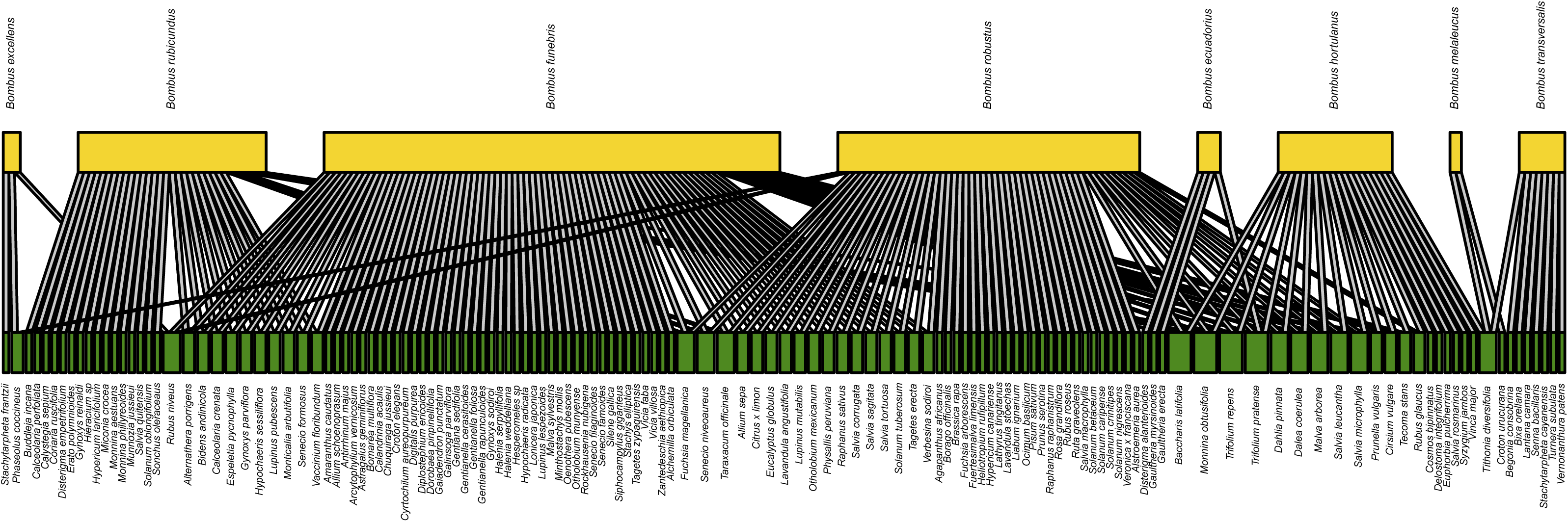
Interaction network between Ecuadorian bumble bees and their associated flowering plants species, created using the bipartite package (Dormann et al., 2014) in R (v.4.4).

**Figure 18.**
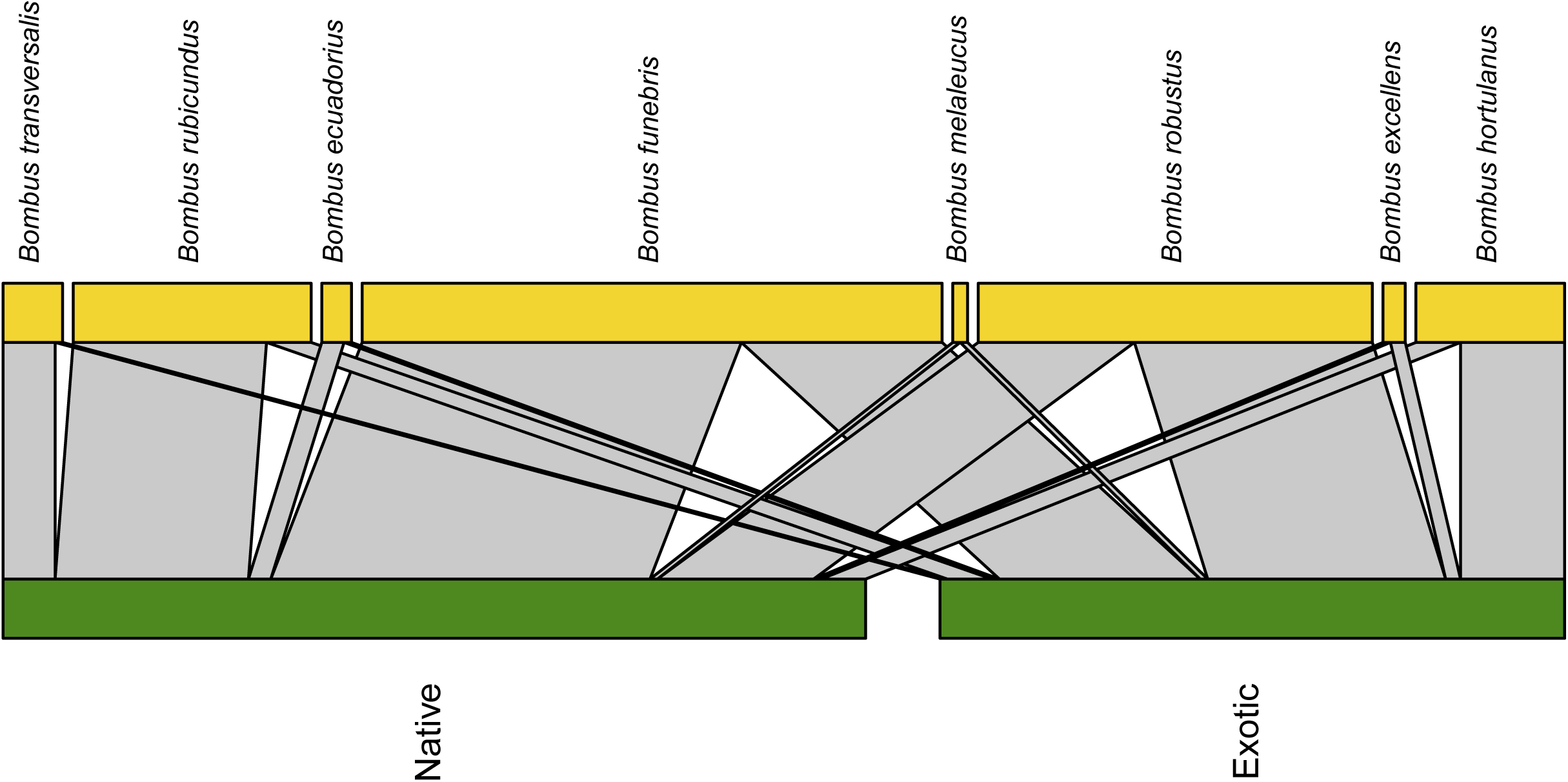
Interaction network between Ecuadorian bumble bees and the origin (native or exotic) of their associated flowering plants species, created using the bipartite package (Dormann et al., 2014) in R (v.4.4).

**Figure 19.**
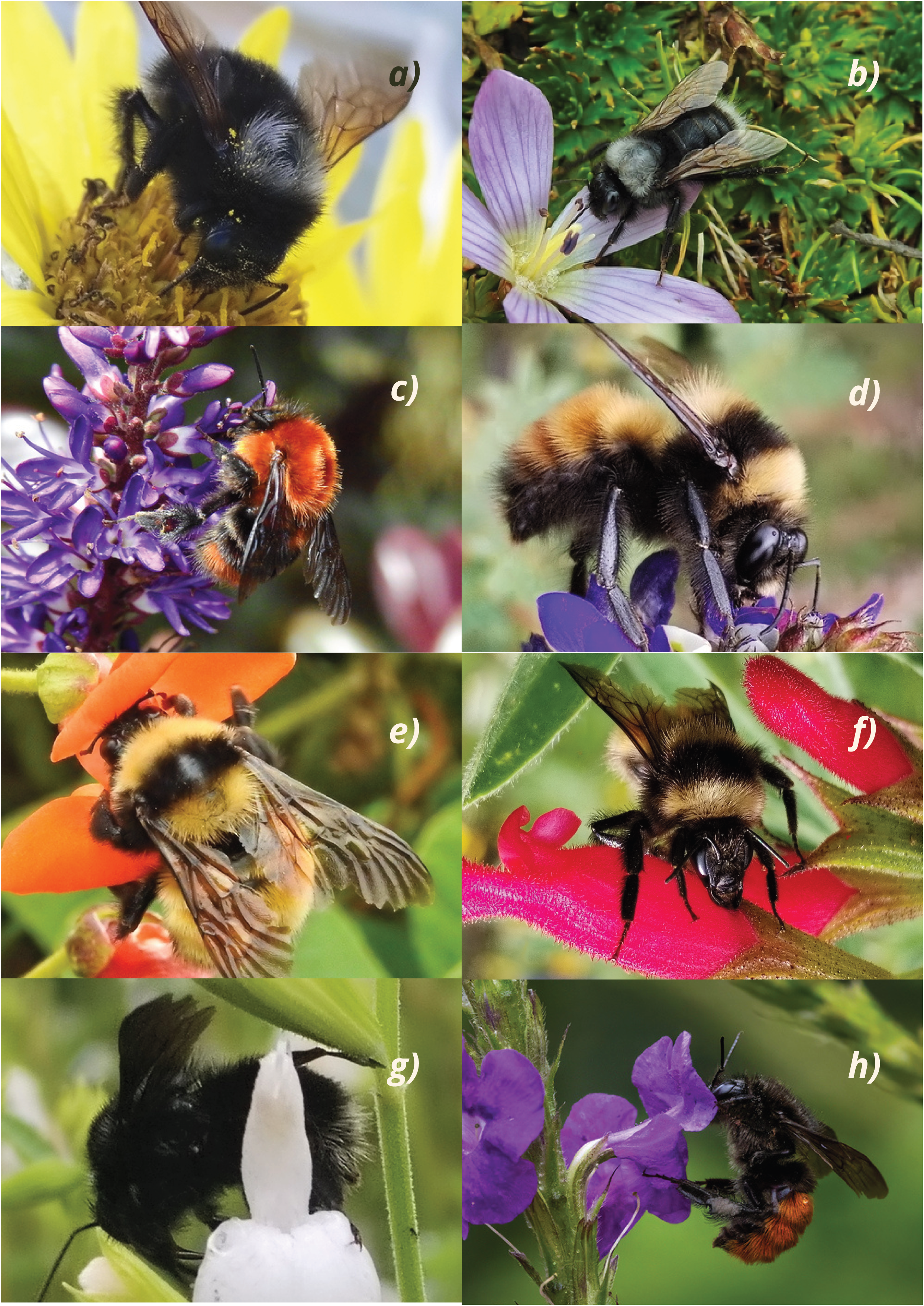
Examples of ecuadorian bumble bees visiting flowers taken from iNaturalist. (a) *Bombus funebris* (bee) and *Gentianella cerastioides* (plant) (credit: Esteban Poveda, user: quilicotriste, iNaturalist: https://www.inaturalist.org/observations/99136154). (b) *Bombus funebris* (bee) and *Senecio niveoaureus* (plant) (credit: Esteban Poveda, user: quilicotriste, iNaturalist: https://www.inaturalist.org/observations/260416732). (c) *Bombus rubicundus* (bee) and *Veronica x franciscana* (plant) (credit: Esteban Poveda, user: quilicotriste, iNaturalist: https://www.inaturalist.org/observations/108821269). (d) *Bombus robustus* (bee) and *Phaseolus coccineus* (plant) (credit: Esteban Poveda, user: quilicotriste, iNaturalist: https://www.inaturalist.org/observations/233124787). (e) *Bombus robustus* (bee) and *Dalea coerulea* (plant) (credit: Esteban Poveda, user: quilicotriste, iNaturalist: https://www.inaturalist.org/observations/297688119). (f) *Bombus robustus* (bee) and *Salvia pauciserrata* (plant) (credit: Esteban Poveda, user: quilicotriste, iNaturalist: https://www.inaturalist.org/observations/274131968). (g) *Bombus vogti* (bee) and *Salvia microphylla* (plant) (credit: Esteban Poveda, user: quilicotriste, iNaturalist: https://www.inaturalist.org/observations/287641794). (h) *Bombus excellens* (bee) and *Stachytarpheta cayennensis* (plant) (credit: Dan Vickers, user: kodiakgarden, iNaturalist: https://www.inaturalist.org/observations/310320696).

The Fabaceae family presented the most floral visits recorded, with 292 records representing 16 species, from which 229 records corresponded to 10 native plant species, and 63 to six exotic plant species. The Asteraceae family presented 137 records from 34 species, from which 94 records corresponded to 25 native plant species and 43 to exotic plant species. The South American native species *Baccharis latifolia* (“chilca”, Asteraceae) and *Monnina obtusifolia* (“iguilán”, Polygalaceae), along with the introduced species *Trifolium repens* (“white clover”, Fabaceae) and *Tithonia diversifolia* (“botón de oro”, Asteraceae), were visited by four bumble bee species: *B. funebris, B. hortulanus, B. rubicundus*, and *B. robustus*, suggesting their role as shared floral resources. The plant species with the highest number of recorded interactions (n = 195) was *Dalea coerulea* (“iso”, Fabaceae), a South American native, with records of visits by *B. funebris, B. hortulanus*, and *B. robustus*.

## Discussion

Our study provides the first catalogue of bumble bees species and reported records of their associated flowering plants in Ecuador, contributing significantly to addressing the regional knowledge gap on bee diversity and distribution. In Ecuador, studies have addressed the taxonomy and distribution of *Bombus* species (González & Engel, 2004; Abrahamovich et al., 2004; Rasmussen, 2003), while others have focused on their natural history (Taylor & Cameron, 2003 studied *B. transversalis*; Nates-Parra, 2006 studied *B. hortulanus*), or on their interactions with flowering plants on natural and agricultural ecosystems (Serpa & Barragán, 2025 and Vanegas & Padrón, 2022 studied *B. funebris)*. We aim to clarify certain aspects of the genus’ distribution and provide updated information regarding its biology.

The exclusion of literature records that corresponded to misattributed collection localities for some bumble bee species clarifies their current distribution and highlights the importance of revisiting historical records localities. Moreover, reports of extensions in altitudinal ranges allow us to better understand their habitat preferences, behavior and nesting biology, especially in highlands species that are adapting to environmental changes in a climate change scenario.

Congruence between morphological and molecular identifications supports the utility of DNA barcoding for rapid biodiversity assessments. Other studies employing DNA barcoding to assess bee diversity have demonstrated that this approach offers a simple and accurate means of associating sexes and castes, many of which are strongly dimorphic and have at times been described as separate species (Sheffield et al., 2009; Janzen et al., 2005). For instance, in the genus *Nomada* (Apidae) and *Sphecodes* (Halictidae) both notoriously difficult to identify reliably given that many species are described from a single sex (Mitchell, 1960, 1962) and exhibit considerable morphological variation (e.g., color patterns, scutellum shape) DNA barcoding has proven effective in linking males and females, thereby enhancing taxonomic descriptions (Sheffield et al., 2009).

In the present study, all barcoded species were assigned to a single BIN, with the exception of *B. hortulanus*, which was divided into two BINs (Supplementary 1, Table 2), likely reflecting genetic divergence between geographically distinct regions such as the Andes and the Ecuadorian Amazonian region. Comparable patterns have been reported in North American bumble bees; for example, in *Bombus ternarius* (Say, 1837), sequence divergence exceeded expected levels (mean = 2.237 ± 0.515% SE; maximum = 4.592%; n = 6). Although this species is among the most easily recognized in eastern Canada (Laverty & Harder, 1988), two genetically distinct groups were identified in Nova Scotia, a pattern further corroborated with the inclusion of additional material from other localities (Sheffield et al., 2003). In Ecuador, expanded barcoding efforts are required to confirm and resolve similar cases.

Citizen science platforms such as iNaturalist proved to be valuable tools for documenting potential plant–pollinator or plant-visitor interactions, as this type of studies usually requires large observation efforts (Beccacece et al., 2025, Bosenbecker et al., 2023). Although records from citizen science platforms tend to be geographically biased, they constitute a useful tool for filling knowledge gaps in understudied interactions (Beccacece et al., 2025). In particular, *Bombus funebris*, a common species, interacted with 80 different plants, while rare species such as *B. ecuadorius*, *B. excellens*, and *B. melaleucus* are represented by few plant interaction records, with a total of 12 records for the three of them (Figure 17, suppl 1). Understanding the population dynamics of both common and rare species is essential for assessing their resilience to environmental change and for designing local conservation strategies. Among plants, the native *Dalea coerulea* (“iso”, Fabaceae) recorded the highest number of visit records in our dataset (n=195), underscoring its role as a keystone resource for bumble bee foraging in the region. Therefore, protecting and promoting native plant species across natural, agricultural, and urban habitats is crucial to enhance floral resource availability and support more resilient pollinator communities.

Given that over half of the documented species are currently listed as Data Deficient by the IUCN (International Union for Conservation of Nature), our findings provide essential baseline data for future assessments. Continued efforts combining natural history, molecular tools, and participatory science are essential for developing conservation strategies and ensuring the long-term persistence of bumble bee diversity in the Tropical Andes.

## Conclusions

This study provides the first comprehensive catalogue of bumble bee species and their associated flowering plants in Ecuador, documenting 15 *Bombus* species across 22 provinces of continental Ecuador, of which eight were recorded interacting with 138 plant species, from which 87 were native. DNA barcoding supported species identification and revealed preliminary patterns of genetic divergence potentially shaped by geographic barriers. Expanding molecular studies, together with natural history data and citizen science, will be key to advancing conservation strategies and safeguarding bumble bee diversity in the Tropical Andes.

## Supporting information

Supplementary 1. Details of museum, literature, GBIF, iNaturalist records, accession numbers and BINs.

## Acknowledgements

We are grateful to the iNaturalist contributors who shared their image-based records and information. We also acknowledge Dra. Priscilla Muriel and Dr. Álvaro Pérez (Herbarium, PUCE) for their support in the identification and validation of plant records. This research was conducted under Research Permit No. MAATE-DBI-CM-2024-0440, issued to the Pontifical Catholic University of Ecuador (PUCE) and MAATE-DBI-CM-2023-0309 issued to INABIO. We thank Nico Blüthgen, Martin Schaefer, María-José Endara, Juan Guevara, Sebastián Escobar, Edith Villa and Karin Römer REASSEMBLY’s project coordination and administration.

